# Towards Atomistic Fidelity in Martini 3: A High-Resolution Protein Model

**DOI:** 10.64898/2026.11.24.766588

**Authors:** Luís Borges-Araújo, Ilias Patmanidis, Valentina Corradi, Kasper B. Pedersen, Thaísa J. Uzan, Hanif M. Khan, Hafez Razmazma, Amita Rani Sahoo, Marc Gueroult, Thilo Duve, Luis F. Cofas-Vargas, F. Emil Thomasen, Fran Bačić Toplek, Christopher Brasnett, Satyajit Mayor, Melanie König, Tugba N. Ozturk, Fabian Grünewald, Solomon Aryee, Kai S. Stroh, Peter C. Kroon, Ana Borges-Araújo, Ayan Majumder, Sangram Prusty, Khaled Mehy-Dine, Sebastian Jimenez-Millan, Alex Peralvarez-Marin, Tatiana I. Morozova, Fabio Lolicato, Alexandre S. de Araujo, Matthieu Chavent, Marco Cecchini, Ivan R. Sasselli, Helgi I. Ingólfsson, Manuel N. Melo, John E. Straub, Matthias Buck, Sebastian Thallmair, Adolfo B. Poma, Phillip J. Stansfeld, Kresten Lindorff-Larsen, Siewert J. Marrink, D. Peter Tieleman, Paulo C. T. Souza

## Abstract

Coarse-grained (CG) protein models enable the simulation of biomolecular processes across extended spatial and temporal scales, but their limited resolution often makes it difficult to capture conformational changes in proteins and peptides in response to their molecular environment. Here, we introduce a higher-resolution Martini 3 protein model featuring a quasi-atomistic backbone representation that maintains numerical stability at conventional Martini timesteps of 20 fs. The model employs a novel bonded scaffold that reconstructs the peptide backbone using virtual sites, enabling a more detailed description of backbone geometry while largely preserving computational efficiency. This architecture allows improved control over backbone torsions and sidechain rotamers, with parameters derived from experimental protein structures and atomistic simulations. Additional refinements to backbone and sidechain non-bonded interactions, together with optional secondary-and tertiary-structure biases and CMAP corrections for disordered proteins, broaden the range of protein and peptide systems that can be represented and the conformational behavior that can be explored. Although the model is not yet capable of reliably reproducing the folding pathways of full proteins, it provides a more flexible representation of proteins and peptides, representing a step toward future foldable Martini protein models.

**Open-beta release and scope:** This preprint accompanies the open-beta release of the high-resolution Martini 3 protein model, publicly available at https://github.com/Martini-Force-Field-Initiative/HighResolutionM3ProteinModel. The release aims to provide early community access to the model, allowing interested researchers to explore its capabilities and incor-porate it into their own simulation workflows ahead of a more comprehensive publication covering its final parameterization, validation, and showcase applications. Both the model and this manuscript remain under active development. While the current implementation is functional and has undergone preliminary validation, several aspects of the parameterization are still being refined, and not all validation results and ongoing developments are presented here. The parameterization and implementation described in this manuscript correspond to the open-beta release `beta_1.0.0`, with subsequent updates and refinements documented through the model repository. We encourage researchers to explore the model, evaluate its performance across different systems, and provide feedback on its accuracy, numerical stability, usability, and potential limitations. Reports of unexpected behavior and suggestions for improvement are welcome through the project’s GitHub repository.

## 1. INTRODUCTION

Coarse-grained (CG) models of proteins are essential tools for exploring biomolecular processes across broad temporal and spatial scales, from local conformational rearrangements to the assembly of large protein complexes and interactions with membranes or ligands[1]. Among CG approaches, the Martini force field [2, 3] stands out for its versatility and widespread adoption, particularly due to the success of its protein model [4]. With the introduction of Martini 3, a refined protein model was also developed [2, 5], building on top of the existing Martini 2 protein model [6, 7] to address key limitations and extend the model’s applicability to a wider range of systems.

In the conventional Martini 3 protein model [2, 5], each residue is represented by a single backbone bead (BB), placed at the atomistic backbone center of mass, to which one or more sidechain beads (SC) are attached. These are connected via harmonic bonds, with additional angle and dihedral potentials applied to sequences of BB beads to reproduce backbone torsional stiffness. Critically, several of these bonded parameters are secondary-structure dependent — assigned based on user-defined structural input — meaning the model cannot fold spontaneously and remains limited to a prescribed conformational landscape. Sidechains are parameterized following Martini 3 guidelines, incorporating molecular volume and partitioning data, and are further stabilized through the sidechain fix (scFix) [8], a torsional restraint mechanism that is itself derived from the initial input structure. To preserve tertiary structure, external biasing schemes such as elastic networks [9] or Gō-like models [5, 10, 11] are required. As a result, despite improvements in sidechain realism and overall transferability, the Martini 3 protein model remains strongly constrained by the input structure at both the secondary and tertiary levels, limiting its ability to capture large-scale conformational flexibility or folding transitions.

Further limitations arise from the coarse resolution of the model itself: with only a single BB bead per residue, the degrees of freedom available for defining backbone torsions and sidechain rotamers are inherently restricted [4]. This minimalistic representation constrains the complexity of bonded potentials that can be used to capture local geometry and conformational variability. For example, backbone dihedrals are limited to generic torsion profiles applied over three or four BB beads, offering little capacity to encode finer structural preferences or local stiffness variations. Similarly, the sidechain conformational landscape can only be crudely modulated, even with scFix corrections, due to the lack of geometric references beyond BB and SC anchor points. While increasing the level of detail in the backbone could, in principle, allow for more accurate structural modeling, doing so often introduces numerical instabilities unless the simulation timestep is drastically reduced. Maintaining stability at Martini’s standard 20 fs timestep becomes a major challenge when adding degrees of freedom. This tradeoff between resolution and numerical performance has been a key barrier to refining backbone architecture.

Despite notable improvements compared to previous iterations, the Martini 3 protein model has displayed several limitations that hinder its broader applicability [4]. Reports have identified issues such as overly compact intrinsically disordered proteins (IDPs) and multidomain proteins (MDPs) [12–14], incorrect membrane insertion of short transmembrane peptides [15–17], weakened protein-protein interactions in solution [11, 17], and excessive solubility of short peptides [18]. While these problems may initially seem unrelated, many can be traced down to two common causes: limited backbone resolution and structure-dependent bonded parameters. The current one-bead-per-residue backbone restricts the complexity of torsional potentials and hampers accurate representation of backbone geometry, particularly in disordered or flexible regions. In coil-assigned residues, insufficient torsional control allows sampling of unphysical conformations, contributing to collapse. Likewise, the inability to shield polar backbone groups and the coarse description of sidechain orientations affect both membrane insertion and protein packing. Together, these issues highlight the need for a higher-resolution, more chemically descriptive backbone model that can decouple structural behavior from user-defined secondary and tertiary restraints.

In this work, we introduce a higher-resolution Martini 3 protein model featuring a quasi-atomistic backbone architecture designed to overcome the limitations of conventional one-bead-per-residue representations. Our approach enables more detailed control over backbone torsions and sidechain rotamers, together with an explicit representation of the backbone dipole, while preserving the hallmark numerical efficiency of Martini through a bonded framework that remains stable at 20 fs timesteps. Although the model is not yet capable of reliably reproducing the folding pathways of full proteins, it offers improved local flexibility and geometric accuracy. Given its ongoing development, optional secondary-structure biases can still be employed when greater conformational stability is required, while also maintaining compatibility with conventional Martini approaches. Overall, these enhancements make the model particularly promising for soft and conformationally dynamic systems, such as intrinsically disordered proteins (IDPs), short peptides, and proteins undergoing structural transitions, where the standard Martini 3 model often falls short. As such, this model represents a step toward future foldable coarse-grained protein models that combine structural plasticity, responsiveness to different molecular environments, and computational efficiency.

## 2. MODEL ARCHITECTURE

### 2.1. Core backbone bonded setup

The high-resolution protein model builds upon the conventional Martini 3 protein model by increasing the granularity of the backbone while preserving its computational efficiency. In the new representation, each amino acid backbone comprises four beads positioned at the corresponding atomistic N, C*α*, C, and O positions, as shown in Figure 1. Increasing the structural resolution of CG protein models is not a novel concept, having been explored to varying extents in other approaches, including UNRES [19], SIRAH [20, 21], PaLaCe [22], and OPEP [23]. Hybrid-resolution approaches such as PACE [24] have also combined detailed protein representations with coarsegrained environments. However, introducing additional backbone detail while retaining the large integration timesteps characteristic of Martini presents a substantial numerical challenge. A central objective of the present development was therefore to identify a bonded arrangement capable of maintaining stability at the conventional Martini timestep of 20 fs. Directly reproducing an atomistic-like bonded topology, with its numerous short bonds, stiff angular potentials, and dihedral terms, would introduce high-frequency motions incompatible with such large integration timesteps.

**Figure 1.**
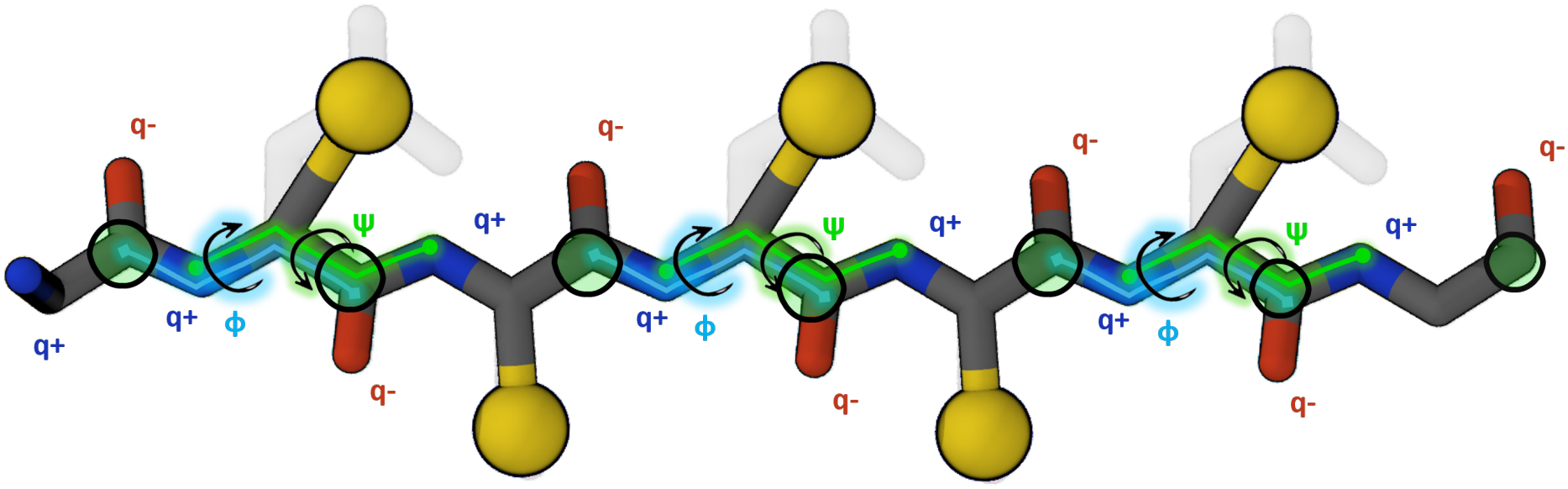
Schematic representation of the high-resolution Martini 3 protein model. The quasi-atomistic backbone comprises four sites corresponding to the N, C*α*, C, and O atoms, each with a distinct role. The carbonyl carbon (C) carries the only backbone Lennard–Jones (LJ) interaction center, represented by an SP1 bead by default. The N and O sites carry partial charges of +0.1*e* and −0.1*e*, respectively, introducing an explicit backbone dipole without additional LJ interactions. The C*α* site serves as a geometric reference for the bonded scaffold and does not participate in non-bonded interactions. C*α* and O are explicitly integrated particles, whereas C and N are reconstructed as massless virtual sites. The triangular bonded scaffold, formed by C*α*_*i*_, O_*i*_, and C*α*_*i*+1_, combines constraints and a harmonic bond to maintain numerical stability at the conventional Martini timestep of 20 fs, while enabling explicit control over backbone *ϕ* and *ψ* torsional preferences. The underlying all-atom structure is shown in a semi-transparent representation to illustrate the correspondence between the atomistic and coarse-grained sites.

To overcome this limitation, we developed a simplified bonded scaffold inspired by previous works on rigid-body Martini cholesterol models [25, 26]. The architecture is built around a triangular arrangement connecting the C*α* and O beads of a given residue to the C*α* bead of the following residue. This triangle provides a stable geometric reference from which the positions of the remaining backbone beads, C and N, are reconstructed as in-plane virtual sites. By defining the peptide backbone geometry through this minimal bonded scaffold, the model avoids the numerous short bonds and stiff angular and dihedral potentials that would otherwise be required to maintain the planarity of the peptide bond. Importantly, this arrangement retains the geometric detail of the quasi-atomistic backbone without introducing additional independently integrated degrees of freedom for the virtual sites.

Our initial implementation employed a fully constrained triangle, as commonly used in conjunction with type-3 virtual site constructions. However, this arrangement resulted in convergence issues with the LINCS constraint solver [27, 28] implemented in GROMACS [29], our primary target MD engine, at the intended integration timestep. To circumvent these numerical limitations, a hybrid bonded/constraint arrangement was adopted. The longest edge of the triangle, connecting consecutive C*α* beads (C*α*_*i*_–C*α*_*i*+1_), was retained as a constraint, taking advantage of the relatively conserved distance between consecutive C*α* atoms across different secondary and tertiary structural motifs. Of the two remaining edges, the longer C*α*_*i*_–O_*i*_ distance was selected for representation by a stiff harmonic bond, while the shorter O_*i*_–C*α*_*i*+1_ distance was retained as a constraint. This choice minimizes the numerical difficulties associated with introducing a stiff harmonic potential along a shorter interparticle distance. This hybrid arrangement substantially improved numerical stability, at the expense of small fluctuations in the position of the O bead and, consequently, in the positions of the C and N virtual sites derived from the triangle. Nevertheless, a fully constrained triangle remains an attractive alternative should future improvements in constraint-solving algorithms permit its stable integration at comparable timesteps.

Finally, the backbone mass is distributed between the two explicitly integrated beads, with masses of 46 and 32 Da assigned to C*α* and O, respectively, while the C and N virtual sites are massless. This results in a total backbone mass of 78 Da per residue, approximately 10% greater than that of the conventional Martini protein backbone. This modest increase in mass further improves numerical stability at large integration timesteps, without substantially affecting the dynamical behavior observed in our preliminary tests.

### 2.2. Backbone non-bonded interactions

Despite the increased backbone resolution, the new model retains a single Lennard-Jones (LJ) interaction center per residue, preserving the overall non-bonded resolution of the conventional Martini 3 protein model. This interaction center is placed at the position of the backbone carbonyl carbon (C), replacing the backbone center-of-mass mapping used in the conventional model.

A key modification concerns the size and chemical identity of this interaction center. In conventional Martini 3, most backbone groups are represented by regular P2 beads, whose relatively large excluded volume can dominate the interactions of residues with smaller sidechains. To better reproduce the molecular volume and packing of the back-bone, we instead adopt a Small (S) bead representation, assigning SP1 as the default backbone bead type. The reduction in bead size is accompanied by an adjustment of the chemical bead type to maintain an appropriate balance of backbone polarity within the Martini 3 interaction matrix.

Residue-specific exceptions were introduced for glycine (SP3), alanine (SP2), and proline (SP1r). The increased polarity and self-interaction strength of glycine account for its greater backbone exposure in the absence of a sidechain. Alanine, whose short sidechain pro-vides limited shielding of the backbone, is assigned the slightly more polar SP2 type. Conversely, proline is represented by an SP1r bead, employing the reduced self-interaction label to account for its distinct amide chemistry and lack of a backbone hydrogen-bond donor.

Beyond the LJ interaction center, the increased backbone resolution also provides an opportunity to introduce directional electrostatic interactions. To this end, partial charges of +0.1*e* and −0.1*e* are assigned to the backbone nitrogen (N) and oxygen (O) sites, respectively. Both are represented by dummy sites, the U-type particles available in Martini 3 for sites that do not contribute LJ interactions. These charged sites are located within the effective excluded volume of the backbone SP1 bead centered at the carbonyl carbon, providing steric shielding that prevents unphysical electrostatic collapse. Charge separation introduces an explicit backbone dipole, providing a simple means of recovering some of the directional character of peptide-backbone interactions that is absent from the conventional single-bead representation.

The model is compatible with the standard Martini 3 non-bonded settings, including the cutoffs for van der Waals and electrostatic interactions and the reaction-field treatment with a relative dielectric constant of 15. The backbone dipoles can also be treated using particlemesh Ewald (PME) electrostatics, although further testing is needed to assess the impact of PME on the behavior of different protein and peptide systems with the new model.

### 2.3. Structural reference dataset

The parameterization of the high-resolution protein model was guided by three complementary structural reference datasets, providing information on backbone geometry, conformational preferences, and sidechain rotamers.

The first consisted of a curated collection of experimentally determined protein structures obtained from the I-TASSER structural database [30]. This dataset provides broad coverage of protein sequences, secondary-structure motifs, and local conformational environments, serving as the primary reference for the parameterization of the general backbone torsional potentials. The selected structures were mapped onto the high-resolution Martini representation, from which distributions of the relevant geometric parameters, including interparticle distances, bond angles, and backbone *ϕ*/*ψ* dihedrals, were extracted.

To complement this structural dataset, atomistic molecular dynamics simulations of ten intrinsically disordered proteins (IDPs) were also employed. All ten proteins were simulated using the CHARMM36m force field [31, 32], while eight were additionally simulated using the DES-Amber force field [33]. These trajectories were mapped onto the high-resolution representation and analyzed following the same procedure as the experimental structures. The resulting distributions provided additional reference data for local backbone geometric parameters that are relatively insensitive to secondary-structure motifs, as well as for selected bonded interactions and corrections specifically developed for disordered proteins.

Finally, residue-specific sidechain conformational preferences were characterized through atomistic simulations of Gly–X–Gly tripeptides, where X represents the amino acid of interest. These systems were simulated using CHARMM and Amber force fields, providing complementary descriptions of sidechain conformational sampling. By isolating each amino acid within a minimal peptide environment, this approach allows the characterization of intrinsic sidechain rotameric preferences while reducing the influence of tertiary interactions and neighboring sidechains. The resulting trajectories were mapped onto the high-resolution representation, and the corresponding sidechain dihedral distributions were extracted to guide the parameterization of the residue-specific rotameric potentials.

Together, these datasets provide complementary structural and dynamical references for the development of the model, combining experimentally observed protein conformations with atomistic sampling of disordered protein ensembles and individual amino acid sidechains.

### 2.4. Backbone torsions

The increased backbone resolution enables a more detailed description of local backbone geometry and conformational preferences. In addition to the triangular bonded scaffold described above, an N–C*α*–C angle potential is introduced to control the geometry between consecutive peptide planes. This angle is relatively conserved across amino acids and secondary-structure motifs, allowing a common bonded potential to be employed throughout most of the protein backbone. Its equilibrium value and force constant were selected to reproduce the corresponding distributions extracted from the structural reference dataset.

The peptide-bond torsion (*ω*) is implicitly fixed by the virtual-site construction, preserving the planar geometry of the peptide group. Given the restricted conformational freedom of the peptide bond, this approximation is reasonable for most protein applications and eliminates an additional degree of freedom that would otherwise require stiff bonded potentials. Nevertheless, the current implementation assumes *trans* peptide bonds and does not account for *cis*/*trans* isomerization [34], a limitation particularly relevant for proline-containing sequences, where cis peptide bonds are more frequently observed.

The remaining backbone conformational degrees of freedom are governed by the *ϕ* and *ψ* dihedral angles. The high-resolution model introduces general backbone torsional potentials designed to accommodate the different conformational states populated by protein backbones. These potentials were parameterized against the overall *ϕ* and *ψ* distributions extracted from the structural reference dataset, encompassing different secondary-structure motifs. Rather than imposing a particular structural state, the resulting potentials aim to reproduce the general conformational preferences of the protein back-bone, allowing individual residues to explore different regions of the Ramachandran landscape.

Importantly, the *ϕ* and *ψ* potentials were fitted independently, targeting their respective one-dimensional distributions rather than explicitly reproducing the full two-dimensional Ramachandran surface. Although correlations between these torsions may emerge from the remaining bonded and non-bonded interactions, they are not directly encoded in the torsional potentials [35]. An additional correction accounting for coupled *ϕ*/*ψ* preferences is introduced for intrinsically disordered proteins, as discussed in a subsequent section.

To account for important residue-specific differences in backbone conformational preferences, separate torsional parameter sets were developed for glycine, proline, and residues preceding proline (preproline). Glycine exhibits increased conformational freedom due to the absence of a sidechain, whereas the cyclic sidechain of pro-line imposes strong restrictions on its backbone geometry. Residues preceding proline also display distinct conformational preferences, motivating their treatment as a separate class. All remaining amino acids share a common set of backbone torsional parameters.

Together, these potentials provide a general description of local backbone conformational preferences without requiring predefined secondary-structure assignments. Nevertheless, additional secondary-structure-dependent torsional potentials can be introduced when the stabilization of particular structural motifs is desired, as described in a following section.

### 2.5. Sidechain setup

The increased backbone resolution also enables a more detailed description of sidechain geometry and conformational preferences. While the sidechain mapping largely follows the conventional Martini 3 protein model, its bonded architecture has been extensively revised to take advantage of the additional geometric reference points provided by the quasi-atomistic backbone. Sidechains are connected to the backbone through a harmonic bond between the C*α* and the first sidechain bead (SC1). Although a constraint would be preferable for several amino acids, the additional constraints already present around the C*α* bead in the backbone scaffold prevent its systematic use without compromising LINCS convergence. A harmonic bond was therefore adopted throughout the model, introducing some flexibility in the backbone–sidechain distance while preserving numerical stability at the target integration timestep.

The availability of additional backbone beads allows sidechain conformational preferences to be controlled through a more detailed arrangement of angle and dihedral potentials. Importantly, all sidechain bonded parameters are now defined exclusively using beads belonging to the same amino acid residue. This contrasts with conventional Martini, where sidechain angles and dihedrals often involve backbone beads from neighboring residues, making their geometries sensitive to the local backbone conformation and, consequently, to the assigned secondary structure. By defining the sidechain bonded terms within each residue, the new architecture allows rotameric preferences to be parameterized independently of the initial protein conformation, with-out requiring the structure-derived sidechain restraints commonly employed in conventional Martini models. All sidechain bonded parameters were reassessed and refined against the reference distributions obtained from the atomistic Gly–X–Gly tripeptide simulations described above. These distributions provide residue-specific references for the local geometry and conformational preferences of individual sidechains, allowing the corresponding bonded potentials to be parameterized without relying on a predefined protein structure.

In parallel with the bonded redesign, selected sidechain non-bonded parameters were also revised. These modifications were motivated in part by the reduction in size of the backbone LJ interaction center, which alters the relative contributions of backbone and sidechain beads to the overall molecular volume and interaction balance. The revisions were further guided by the partitioning behavior of sidechain analogues, including octanol–water partition coefficients (LogP), membrane translocation potentials of mean force (PMFs), and the Wimley–White peptide hydrophobicity scale[36]. Observations from previous applications of the Martini 3 protein model, particularly the insufficient membrane affinity of certain transmembrane peptides, also motivated the reassessment of selected sidechain interactions. Among the hydrophobic amino acids, alanine and valine were reassigned from TC3 to TC2 and from SC3 to SC2, respectively. Isoleucine and leucine were both reassigned from SC2 to C1r, maintaining reduced self-interactions while increasing their overall hydrophobicity. Additional refinements were introduced for selected polar and chemically distinct amino acids: asparagine was reassigned from SP5 to P5, cysteine from TC6 to SC6 and the first sidechain bead of arginine from SC3 to SC4. Together, these modifications provide a more detailed and transferable description of sidechain geometry and conformational preferences, while rebalancing sidechain interactions to account for the redesigned backbone representation. While some of these sidechain refinements may also benefit the conventional Martini 3 protein model, their parameterization was performed in the context of the new high-resolution backbone representation. Their transfer-ability to the conventional model therefore remains to be evaluated.

#### 2.5.1. D-amino acids

The increased backbone resolution also enables an explicit representation of amino-acid stereochemistry, allowing the model to distinguish between L-and D-amino acids. Unlike conventional Martini, where the limited backbone resolution prevents an explicit description of backbone chirality, the new architecture provides sufficient geometric reference points to define the stereochemical configuration of individual residues. In the current implementation, D-amino acids are obtained by inverting the equilibrium geometry of the SC1–C*α*–N–C dihedral, while retaining the SC1–C*α*–O–C dihedral unchanged. The backbone *ϕ* and *ψ* torsional potentials are also mirrored to account for the inverted conformational preferences of D-amino acids in Ramachandran space. No additional parameterization was performed for D-amino acids, with the remaining bonded and non-bonded parameters inherited from their corresponding L-amino acid counterparts. Future refinements may further improve their conformational description, particularly in mixed L/D sequences, where neighboring residues of opposite chirality may give rise to distinct local structural preferences.

### 2.6. Terminal modifications

To accommodate different peptide and protein terminal chemistries, the high-resolution model provides several options for N-and C-terminal modifications. The available N-terminal variants include the standard protonated amine (N-ter), the neutral amine (NH2-ter), and the acetylated N-terminus (NACE-ter). For the C-terminus, the model supports the standard deprotonated carboxylate (C-ter), the protonated carboxylic acid (COOH-ter), the methyl ester (CT1-ter), the primary amide (CT2-ter), and the N-methylamide (CT3-ter). These terminal variants were developed using the corresponding atomistic representations available in CHARMM as structural references.

### 2.7. Secondary-and tertiary-structure bias

Although the general backbone torsional potentials allow the model to sample different local conformations without predefined secondary-structure assignments, additional structural biases can be introduced when the preservation of specific secondary or tertiary structural motifs is desired.

#### Secondary-structure bias

Secondary-structure-dependent backbone torsional potentials were developed following the same parameterization approach described for the general *ϕ* and *ψ* dihedrals. However, rather than considering the overall conformational distributions, the structural reference dataset was classified according to the secondary-structure assignments obtained using DSSP [37]. The corresponding *ϕ* and *ψ* distributions were then extracted separately for each structural motif and used to parameterize the respective torsional potentials. As in conventional Martini, the model supports five secondary-structure classes: coil, *α*-helix, *β*-strand, bend, and turn. For coil and *α*-helical conformations, separate torsional parameter sets were developed for glycine, proline, and residues preceding proline (pre-proline), in addition to the general amino acid class. For the remaining secondary-structure motifs, a single set of torsional parameters is employed for all residues. These potentials provide an additional level of control over backbone conformational preferences, allowing specific structural motifs to be stabilized when required. Importantly, unlike the conventional Martini protein model, the underlying backbone architecture (such as bonds, or angle potentials) does not depend on these assignments.

#### Tertiary-structure bias

Although the increased backbone resolution and refined bonded potentials provide improved control over local protein geometry, the model is not yet capable of reliably reproducing the folding pathways of full proteins or maintaining arbitrary folded structures without additional structural biases. This limitation may partly reflect the simplified description of directional hydrogen bonding and single-water-mediated interactions, which play important roles in protein folding and structural stability. For tertiary-structure stabilization, three complementary approaches are currently supported: elastic networks [9], Gō-like models [5], and OLIVES [11]. Further details on the implementation and parameterization of these structural biases will be provided in a future update of this preprint.

### 2.8. CMAP corrections for disordered proteins

While the general backbone torsional potentials provide a description of the individual *ϕ* and *ψ* conformational preferences, their independent parameterization does not explicitly account for correlations between the two dihedral angles [35]. This limitation is particularly relevant for intrinsically disordered proteins (IDPs) and intrinsically disordered regions (IDRs), where the absence of persistent structural constraints makes the sampled conformational ensemble more sensitive to the underlying backbone torsional preferences. Inaccuracies in these preferences can therefore propagate into global ensemble properties, including protein dimensions. The importance of conformational preferences for IDP ensembles was also highlighted in the development of Martini3-IDP, where refinements to backbone and sidechain bonded interactions reduced excessive protein compaction without requiring a general rescaling of non-bonded interactions [14]. To address the missing correlations between backbone torsions, we introduce an additional correction based on the correction map (CMAP) formalism. CMAP potentials provide a two-dimensional energetic correction as a function of the backbone *ϕ* and *ψ* dihedral angles, allowing direct modulation of the coupled conformational landscape [35]. In the current high-resolution Martini 3 model, this correction is applied on top of its coil-specific backbone torsional potentials, improving the description of local conformational preferences in disordered regions without imposing additional structural biases. An equivalent correction for the general backbone torsional potentials, aimed at improving conformational sampling across different secondary-structure motifs, is currently under development.

The correction was developed using the atomistic IDP reference dataset described above. As a starting point, the corresponding proteins were simulated using the high-resolution Martini model with the general backbone torsional parameters, without CMAP corrections, and the resulting *ϕ*/*ψ* distributions were compared against those obtained from the atomistic simulations. The differences between the atomistic and CG Ramachandran distributions were used to construct an initial two-dimensional correction potential, favoring regions underpopulated by the CG model and penalizing those that were excessively sampled. The resulting potential was subsequently smoothed to avoid sharp energetic variations between neighboring regions of the map, which could compromise numerical stability at the target integration timestep. Additional manual adjustments were introduced in selected regions of the Ramachandran landscape to further improve conformational sampling and numerical stability. The resulting CMAP correction was then evaluated through additional simulations of the reference IDP systems. The introduction of the correction improved the conformational behavior of disordered proteins, particularly in longer sequences where small differences in local backbone preferences can accumulate and substantially affect global protein dimensions. The CMAP correction is therefore currently recommended for simulations of IDPs and IDRs using the coil-specific backbone torsional parameters, particularly in the absence of additional secondary-or tertiary-structure biases.

### 2.9. Tool compatibility

To facilitate adoption and integration into existing Martini workflows, the high-resolution protein model has been implemented in Martinize2 [38] and Polyply [39]. These tools provide support for the generation of protein topologies and the construction of molecular systems using the new backbone representation, allowing users to prepare simulations through familiar Martini workflows. Although the high-resolution model is compatible with the broader Martini ecosystem, direct conversion of existing systems using conventional Martini protein representations is not yet supported by the current tools. Dedicated tools to facilitate conversion between Martini protein representations are therefore being considered for future development. Support for the new representation is also being considered for backmapping tools, including CG2AT2 [40] and ezAlign [41], with the aim of enabling conversion between the high-resolution Martini model and atomistic representations. This will facilitate the integration of the model into multiscale simulation workflows, including the refinement and atomistic validation of conformations sampled at the coarse-grained level.

## 3. PRELIMINARY RESULTS AND OUTLOOK

Preliminary tests of the high-resolution Martini 3 protein model suggest improvements in several areas where limitations of the conventional Martini 3 protein model have previously been reported[4]. In particular, the revised backbone and sidechain representations appear to provide a better balance between conformational flexibility and molecular interactions, improving the description of intrinsically disordered proteins, peptide–membrane insertion, and protein–lipid interactions in the systems examined.

The improved conformational behavior also permits a reduction in secondary-and tertiary-structure biases while maintaining the stability of folded proteins in the tested systems. Beyond addressing existing limitations, the increased backbone resolution opens new possibilities, including environment-dependent conformational transitions in short peptides, multistate protein models, and ion permeation through narrow protein channels. However, these capabilities remain system-dependent, and further refinement of the default parameters is needed to achieve a better balance between secondary-structure stability and conformational flexibility across different proteins and peptides.

Despite these encouraging results, the model remains under active development, and further optimization and validation are needed. One outstanding challenge concerns achieving a transferable balance of protein–lipid interactions. While preliminary tests have shown improvements for some peripheral proteins, particularly in their interactions with specific lipids, other systems still exhibit discrepancies with reference data. Results for transmembrane proteins have also been mixed, with some lipid interaction patterns showing good agreement with atomistic simulations and others requiring further investigation. These differences may also depend on the lipid models employed, particularly given recent refinements to the Martini 3 lipidome [42]. Improving this balance of protein–lipid interactions, alongside further assessment of protein–protein interactions, is an active focus of our ongoing work. Nevertheless, the current developments represent a step toward more flexible and transferable Martini protein models while preserving Martini 3’s computational efficiency and compatibility with established simulation workflows.

A more comprehensive assessment of the model, including additional benchmarks and applications, will be presented in an updated version of this preprint and in the final publication.

## 4. ACKNOWLEDGEMENTS

The authors would like to thank Roberto Covino and Romuald Marin for their contributions to the early exploration of secondary-structure-dependent dihedral potentials during the initial stages of model development. We also thank Amauri Donadon, Çağla Okyay, Nitin Kumar Singh, Tristan Bereau, Luis J. Walter, Liguo Wang, Tania Rajpersaud, Yang Liu, Sharon M. Loverde, Giulia Rossetti, Simone Albani, Anna Duncan, Attilio Vittorio Vargiu, Herre Jelger Risselada, Riccardo Alessandri, Leonhard Josef Starke, Rita A. Roessner, Nicolas Floquet, and Theo Cardozo Brascher for their contributions through preliminary testing, scientific discussions, and valuable feedback at different stages of model development.

L.B.A. and P.C.T.S. acknowledge the support of the French National Centre for Scientific Research (CNRS). L.B.A. and S.A. acknowledge support from the Agence Nationale de la Recherche (ANR, France) through the CoarseScreen project (ANR-25-CE11-7960-01). T.J.U. and A.S.A. acknowledge financial support from the Brazilian National Council for Scientific and Technological Development (CNPq, Process No. 404205/2024-0) and the São Paulo Research Foundation (FAPESP, Process No. 2025/23025-6). T.I.M. acknowledges support by the French National Research Agency (Agence Nationale de la Recherche, ANR) through the project DiPCaM, project number ANR-25-CE06-0062-01. P.C.T.S. acknowledges support through research collaboration agreements with Sanofi and PharmCADD. This work was granted access to the HPC resources of IDRIS, CINES, and TGCC under the allocations 2023-A0150314644, 2024-A0160713456, 2025-A010713456, 2026-A0180713456 and 2027-A0200713456 made by GENCI. S.A., L.B.A., T.J.U., and P.C.T.S. acknowledge the support of the PSMN (Pôle Scientifique de Modélisation Numérique) and the Centre Blaise Pascal’s IT test platform at ENS de Lyon (Lyon, France) for computational resources. The platform operates the SIDUS solution developed by Emmanuel Quemener [43].

Work in D.P.T.’s group was supported by the Natural Sciences and Engineering Research Council of Canada and the Canadian Institutes of Health Research. Further support came from the Canada Research Chairs program. Calculations were supported by the Digital Research Alliance of Canada. T.N.O. and H.I.I. thank LDRD 26-SI-004 and 26-ERD-027 for funding and the Livermore Institutional Grand Challenge for computing time. Part of this work was performed under the auspices of the U.S. Department of Energy by Lawrence Livermore National Laboratory under Contract DE-AC52-07NA27344. LLNL-JRNL-2026979. S.J.M. is supported by funding from the European Research Council (ERC) with the Advanced grant 101053661 “COMP-O-CELL”. A.R.S. and M.B. were partially funded by NIH grants R01AG089561 and previously R01EY029169 and we want to thank CWRU HPC for providing computing resources. T.D. and S.T. gratefully acknowledge the computing time provided at the NHR Center NHR@SW at Goethe University Frankfurt (project number 26814). F.G. acknowledges funding from the Giersch Foundation. This work was supported by the Deutsche Forschungsgemeinschaft to FL (SFB-1638/1 - 511488495 - Z01; SFB/TRR 186, project A1; FL, DFG LO 2821/1-1; FL). F.L. acknowledged the computing resources provided by the state of Baden-Württemberg through bwHPC and the German Research Foundation (DFG) through grant INST 35/1597-1 FUGG. F.L. gratefully acknowledge the data storage service SDS@hd supported by the Ministry of Science, Research and the Arts Baden-Württemberg (MWK) and the German Research Foundation (DFG) through grant INST 35/1503-1 FUGG. This work was funded by the Deutsche Forschungsgemeinschaft (DFG, German Research Foundation) under Germany’s Excellence Strategy – EXC-3094 – 533751785 (to S.T.), the Hessian Ministry of Science and Arts LOEWE program (Center for Multiscale Modeling in Life Sciences, CMMS) (to S.T.), the Alfons und Gertrud Kassel Foundation (to S.T.), the Dr. Rolf M. Schwiete Foundation (to S.T.), and the H. & E. Kleber Foundation (to T.D. and S.T.).

The authors used generative artificial intelligence tools to assist with language editing and improving the clarity of the manuscript. All generated content was critically reviewed and revised by the authors, who take full responsibility for the scientific content and conclusions presented.

## Notes

https://github.com/Martini-Force-Field-Initiative/HighResolutionM3ProteinModel

### Competing Interest Statement

K.L.-L. holds stock options in and is a consultant for Peptone. The remaining authors declare no competing interests.

